# Drug-Target Interaction prediction using Multi-Graph Regularized Deep Matrix Factorization

**DOI:** 10.1101/774539

**Authors:** Aanchal Mongia, Angshul Majumdar

## Abstract

Drug discovery is an important field in the pharmaceutical industry with one of its crucial chemogenomic process being drug-target interaction prediction. This interaction determination is expensive and laborious, which brings the need for alternative computational approaches which could help reduce the search space for biological experiments. This paper proposes a novel framework for drug-target interaction (DTI) prediction: Multi-Graph Regularized Deep Matrix Factorization (MGRDMF). The proposed method, motivated by the success of deep learning, finds a low-rank solution which is structured by the proximities of drugs and targets (drug similarities and target similarities) using deep matrix factorization. Deep matrix factorization is capable of learning deep representations of drugs and targets for interaction prediction. It is an established fact that drug and target similarities incorporation preserves the local geometries of the data in original space and learns the data manifold better. However, there is no literature on which the type of similarity matrix (apart from the standard biological chemical structure similarity for drugs and genomic sequence similarity for targets) could best help in DTI prediction. Therefore, we attempt to take into account various types of similarities between drugs/targets as multiple graph Laplacian regularization terms which take into account the neighborhood information between drugs/targets. This is the first work which has leveraged multiple similarity/neighborhood information into the deep learning framework for drug-target interaction prediction. The cross-validation results on four benchmark data sets validate the efficacy of the proposed algorithm by outperforming shallow state-of-the-art computational methods on the grounds of AUPR and AUC.

## Introduction

Genomic drug discovery is one of the key branches of Pharmaceutical Sciences. The task in drug discovery is to search for adequate interactions between targets (proteins or amino acid sequences) and drugs (chemical compounds). Conventionally, this was done through time-taking and expensive wet-lab experiments. In recent times, the introduction of computational techniques for prediction of interaction probability [1–4] has paved the way for appropriate and effective alternatives which could help avoid costly candidate failures. These methods take some existing experimentally valid interactions which are publicly available in databases like STITCH [5], ChEMBL [6], KEGG DRUG [7], DrugBank [8] and SuperTarget [9] to predict the interaction probability of unknown drug-target pairs. Successfully identifying the compound-target interaction not only assists drug discovery but also affects other fields such as drug repositioning, drug resistance and side-effect prediction [10]. As an example, Drug repositioning [11,12] (using an existing drug for new indications) can grant polypharmacology (multi-target effect) to a drug. Gleevec (imatinib mesylate) is one of the many such examples which was successfully repositioned. Earlier, it was known to interact only with the Bcr-Abl fusion gene which is indicative of leukemia. However, later discoveries showing that it also interacts with PDGF and KIT, repositioned it for the treatment of gastrointestinal stromal tumors [13,14].

The methods available for prediction of DTI can be divided into the following three broad categories: Ligand-based approaches, Docking based approaches, and Chemogenomic approaches. Ligand-based approaches predict interactions by deploying the similarity between the ligands of target proteins [15]. The idea is that molecules with similar structure/property would bind similar proteins [16]. But, the reliability of results might get compromised due to limited information about known ligands per protein. Docking-based approaches use the three-dimensional structure of both drugs and proteins for prediction of the interaction probability/likelihood [17–19]. This, although is well-accepted, but is very time-taking and hence, is not a feasible solution for some families of proteins for which the 3D structure is unavailable or is difficult to predict [20] like in case of the GPCRs (G-protein coupled receptors).

The ability of Chemogenomic approaches to overcome the challenges faced by conventional methods have made them quite popular in the past few years. Such approaches can process huge amount of biological databases, existing publicly on the web and can process metadata (chemical structures and genomic sequences) for both the drug and target, respectively. These methods can be further be divided on the basis of input data representation: Feature-based techniques and Similarity-based techniques. Feature-based techniques take their inputs in the form of features and class labels (binary values here) and leverage machine learning for classifying if an input instance corresponds to a positive interaction or a negative one. For example, Decision Trees (DT), Random Forests (RF) [25], ensemble-based technique [21] and Support Vector Machines (SVM). In the training set, positive samples are the experimentally known interactions while the negative ones are either non-interactions or unknown interactions. The second category includes Similarity-based methods, which take into account two similarity matrices corresponding to drug and target similarity, respectively, along with the drug-target interaction matrix.

Amongst the recent techniques proposed for DTI prediction, matrix factorization based models depict the most promising results [2]. There has been recent work on deep Matrix Factorization [22] too. [22] proposes a deep neural network based model with Matrix Factorization based learning. However, none of them graph regularize the metadata associated with the drugs and targets (similarity information). Amongst the shallow techniques in the literature, graph regularized version of Nuclear norm minimization [23] and matrix factorization [24] have been proposed to incorporate drug and target similarities. But, these approaches solve the respective cost function by regularizing it using the standard similarity for drugs (*S*_*d*_) and targets (*S*_*t*_). These standard similarities have been computed from the chemical structure of drugs using SIMCOMP score and from target genomic sequences using Normalized Smith-Waterman score respectively [25]. These proposed methods do not explore the multi-modality of the metadata. The similarities could be computed using other metrics like cosine similarity, Jaccard similarity, etc. To overcome this, multiple graph laplacian terms for both drugs and targets have been augmented to deep matrix factorization cost function in order to incorporate multiple types of similarities between the drugs as well as the targets. In other words, we have taken advantage of learning complex deep representations as well as incorporating multiple similarities to predict the Drug-Target interactions by graph regularizing deep matrix factorization with multiple similarities.

The Graph Laplacians are constructed using the standard similarity measures available for drugs (chemical structure similarity) and targets (genomic sequence similarity) and four new kinds of similarities which are computed from the DTI matrix itself. These include the Cosine similarity, Correlation, Hamming distance and Jaccard similarity between the drugs/targets. There exists no prior work in DTI prediction which uses Deep Matrix Factorization regularized by multiple graph laplacians.

## Background

The aforementioned DTI problem and the problem of collaborative filtering (CF) has a lot in common. In information retrieval, CF is a standard problem which is used in recommendation systems (e.g. in Amazon product recommendations, Netflix movie recommendations, etc). It is built upon a dataset of users and items/movies which they rate. Since the number of items are humongous and not all of these can be rated by the users, the rating dataset is sparse. Therefore, the objective is to estimate the ratings of all the users for all the items. Correct estimation eventually assists in recommending new items/movies to other users and improves prediction accuracy. In DTI, the drugs play the role of users, targets can be thought of as items, and interaction matrices are similar to rating matrix. In the past few years, a lot of techniques used for DTI prediction were in fact, originally developed for CF.

The initial techniques developed in both the fields (CF [26] and DTI [27–29]) were simple and based on neighborhood-based models. To predict the interaction of an active drug and target, firstly similar or neighboring drugs are identified (using some notion of similarity). This is done in order to weigh (by the normalized similarity score) the neighborhood’s interaction for predicting the interaction of an active drug/target. The second set of approaches were based on bipartite local models where a local model is built for each drug and target. For instance, in [30], an SVM is trained to predict the interaction of each drug on all targets and each target on all drugs. The prediction provided by the model in both cases is merged to generate the final interactions. Many other techniques such as [31–33] fall under this generic approach.

The third approach is based on network diffusion models. To predict DTI [34] used a model which depends on a random walk of the network with a predefined transition matrix. Another approach based on network diffusion model finds a simple path (without loops) between different network nodes for predicting interactions.

The fourth approach, matrix factorized based prediction, again was initially implemented for collaborative filtering [35,36]. Matrix factorization assumes that the data matrix can be factorized into two latent factor matrices, one corresponding to drugs and another to targets. If the latent factors of the drug and the target match (i.e. when the inner product has a high value) then this results in a high probability of their interaction. Hence, the ability to express the interaction matrix as the product of the two latent factor matrices enables matrix factorization along with its variant to be useful for solving the DTI problem [24,37].

Lastly, feature-based or classification based approaches were also proposed, where the drugs and targets are represented as features, using their chemical or biological data. Varied set of feature selection methods [38,39] can be used for the same. These features are further concatenated, while the interaction associated with the concerned drug-target pair is assigned as the label for this entire feature; later any standard classifier is used for prediction of label/interaction.

In one of the recent studies [2], it was shown that matrix factorization works best for DTI prediction. This method assumes that there are very few (latent) factors which are actually responsible for drug target interactions. So, DTI matrix can be factored into a tall (drug) latent factor matrix and a fat (target) latent factor matrix. Mathematically, the assumption is that the DTI matrix is of low-rank. Matrix factorization is being used to model low-rank matrices for the past two decades since the publication of Lee and Seung’s seminal paper [40].

### Matrix Factorization

Matrix factorization has previously been used to find low dimension decompositions of matrices in a number of applications [41,42]. It is one of the direct methods to solve the low-rank matrix completion problem. Let us assume that *X*_*m*×*n*_ (known to have rank *r*) is the adjacency matrix where each entry denotes interaction between a drug and target (1 if they interact, 0 otherwise). Unfortunately, we only observe this matrix partially because all interactions are not known. If *Y* denotes the partially observed adjacency matrix, the mathematical relation between *X* and *Y* is expressed as:

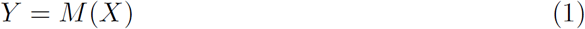

In the above equation, *M* denotes the binary mask which has 1’s where the interactions have been observed and 0’s where they are unobserved or not known. It is also called the sub-sampling operator. The problem is to find *X*, given the sparse observations *Y* and sub-sampling operator *M*. *X*_*m*×*n*_ can be expressed as a product of two matrices *U*_*m*×*r*_ and *V*_*r*×*n*_. The complete matrix factorization problem (1) can be framed as, (2).

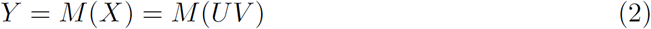

Predicting *U* and *V* from (2) tantamount to estimating X. These two-factor matrices (*U* and *V*) can be solved by minimizing the *Frobenius* norm of the following objective function.

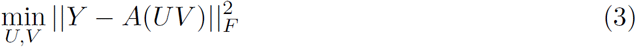

The global convergence is not guaranteed, given the bi-linear nature of the problem; but it usually works in practice.

### Deep Matrix Factorization

In recent times, deep learning has permeated almost every aspect of computational science, including Bioinformatics and Drug discovery [43–45]. Our current work is motivated by the success of deep matrix factorization [46–48] and deep dictionary learning [49]. The basic idea in there is to factor the data matrix into several layers of basis and a final layer of coefficients; shown here for two levels –

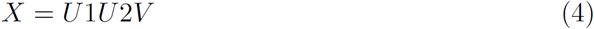

For visual understanding, please refer to the decomposition shown in Figure 1. Note that this is a feed backward neural network, the connections are from the nodes towards the input. This is because matrix factorization is a synthesis formulation. Incorporating the deep matrix factorization formulation into (2) leads to,

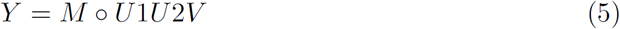

Our task is to solve the different layers of basis (U1, U2) and the coefficients (V) by solving the least squares objective function.

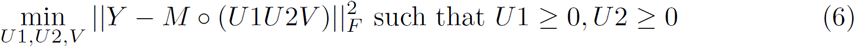

**Fig 1.** Multi-Graph regularized Deep Matrix Factorization pipeline.

## Proposed Formulation: Multi-Graph Regularized Deep Matrix Factorization

In the past studies, it has been shown that taking metadata or similarity information for drugs/targets into account greatly improves the DTI prediction [25]. But, the standard Matrix Factorization and Deep Matrix Factorization algorithms cannot accommodate such metadata. There are a couple of recent works which do use standard similarities for drugs and targets with matrix factorization [24,50] and matrix completion [23] frameworks. It is imperative that DMF should be able to take into account the standard similarity information as well as more types and combinations of similarities. To accomplish this, four other types of similarities (between drugs/targets) have been augmented to presented Multi-Graph regularized Deep Matrix Factorization (MGRDMF).

Multigraph regularization is incorporated into the objective function of Deep matrix factorization by taking into account the Laplacian weights corresponding to drugs and targets. The cost function for a multi-graph regularized 2-Layer deep matrix factorization is given by:

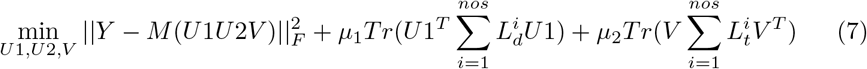

where *μ*_1_ ≥ 0 and *μ*_2_ ≥ 0 are parameters to penalize the regularization terms, *Tr*(.) is the trace of a matrix, *nos* denotes the number of similarity matrices (*nos* = 5 here).

If, say we consider a single similarity matrix for drugs (*S*_*d*_) and that for targets (*S*_*t*_), then *L*_*d*_ = *D*_*d*_ − *S*_*d*_ and *L*_*t*_ = *D*_*t*_ − *S*_*t*_ are the graph Laplacians [51] for *S*_*d*_ (drug similarity matrix) and *S*_*t*_ (target similarity matrix), respectively, and 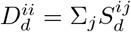 and 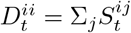 are degree matrices.

Problem (7) is soved using Majorization-Minimization [52] to de-couple the mask *M* transforming the objective function to:

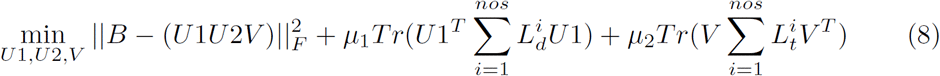

where 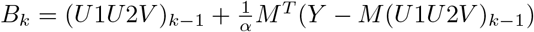. Here, *k* denotes the iteration count. To find a solution to the above problem, we need to solve for variables *U*1, *U*2 and *V*:

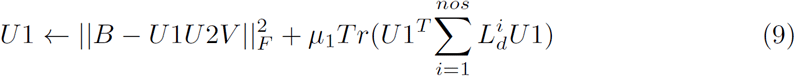

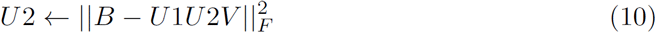

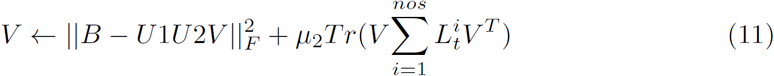

For solving *U*2, it can be iteratively updated as follows:

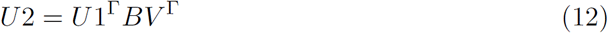

To solve for *U*1, we differentiate eq (9) wrt *U*1 and equate it to 0:

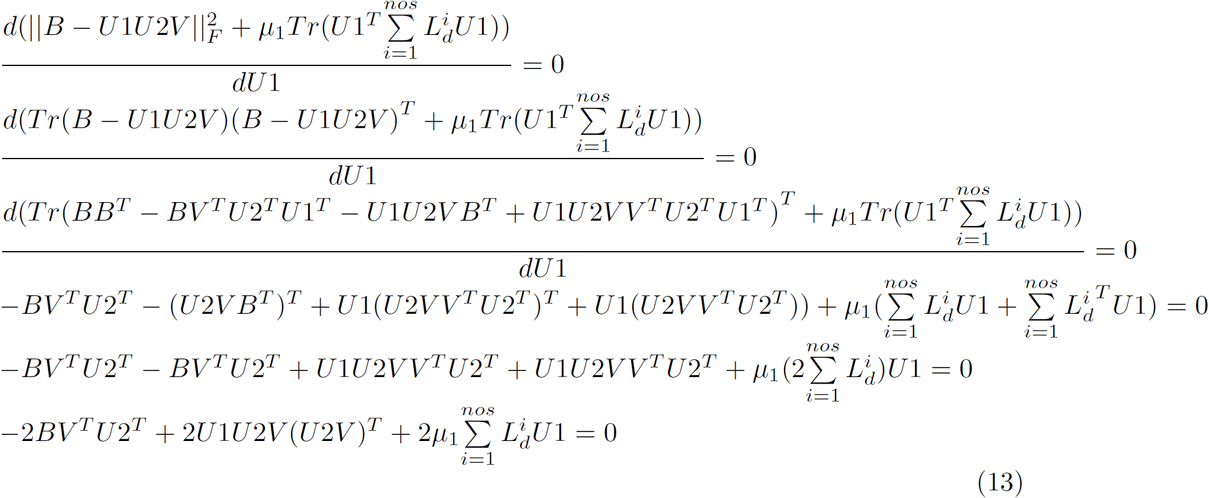

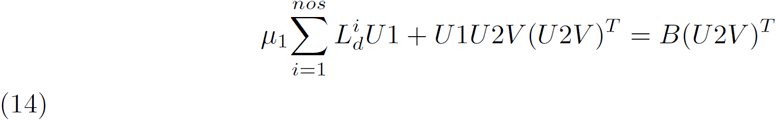

This is the form of a sylvester equation, an equation of the form: *A*_1_*X* + *XA*_2_ = *A*_3_ (Here, 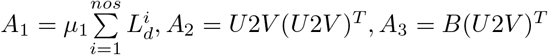). It has a unique solution if the eigen values of 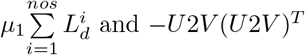 and −*U*2*V*(*U*2*V*)^*T*^ are distinct.

A similar derivation can be followed to solve for V to get the following sylvester equation:

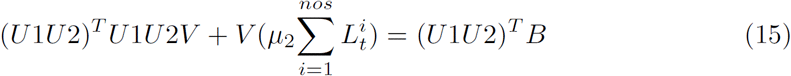

One can argue that computing the Laplacian using the similarity matrix which is obtained by summing up various similarity matrices 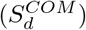 is nothing but the sum of all the individual Graph Laplacians involved. The mathematical proof for combined graph Laplacian has been shown below:

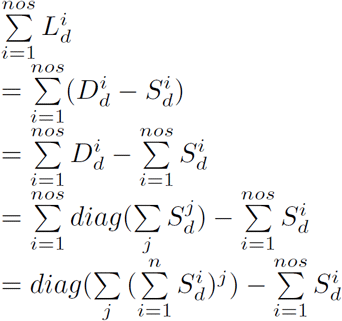

Let 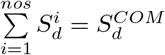 where 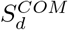 denotes combined similarity for drugs. Essentially,

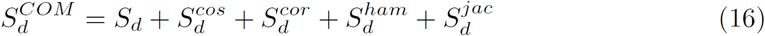

ł Then,

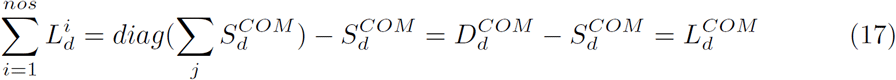

Here, 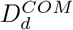 and 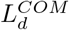 stands for combined degree matrix and combined Laplacian matrix (summation of all graph laplacians) for drugs. The pseudo-code for the method has been shown in Algorithm 1.

### Algorithm 1 Multi Graph regularized Deep Matrix Factorization

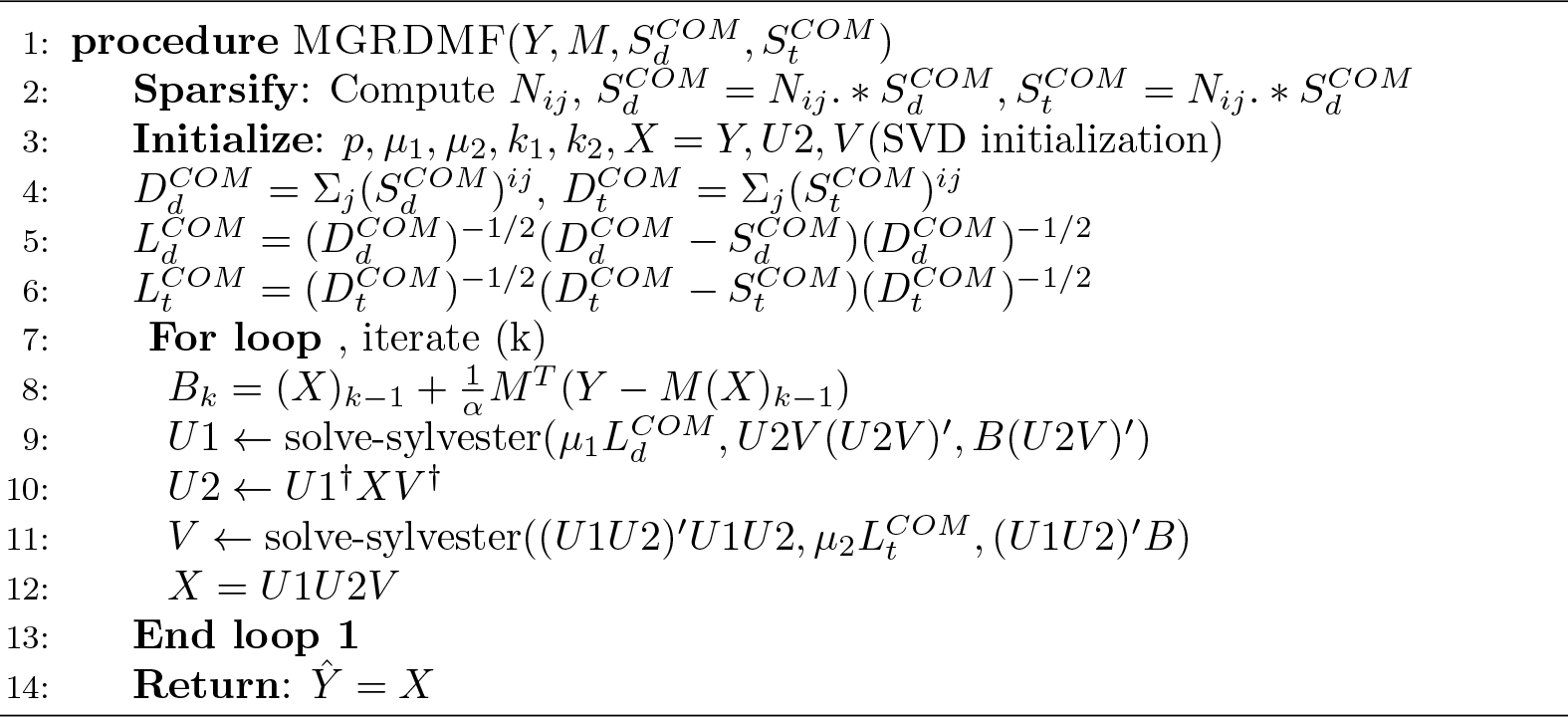

## Results and Discussion

### Dataset Description

To evaluate the performance of the proposed technique and the baseline methods, we use four types of drug-target interaction data (concerning 4 protein classes-E: enzymes, IC: ion channels, GPCR: G protein- coupled receptors and NR: nuclear receptors) provided by [27]. Along with the interactions, associated compound-structure/drug similarity, *S*_*d*_ (measured using SIMCOMP [53]) and protein-sequence/target similarity, *S*_*t*_ (computed using normalized Smith–Waterman [54]) are also provided. The interaction information is publicly available at http://web.kuicr.kyoto-u.ac.jp/supp/yoshi/drugtarget/ and has been obtained from public databases such as KEGG BRITE [55], BRENDA [56] SuperTarget [9] and DrugBank [8].

This data is modelled as an adjacency matrix between drugs and targets, and encodes the interaction between *n* drugs and *m* targets as 1 if the drug *d*_*i*_ and target *t*_*j*_ are known to interact and 0, otherwise.

The similarity matrices *S*_*d*_ and *S*_*t*_ constitute the most standard similarities that have been used in the DTI prediction task hitherto. We use these similarities along with the following four more similarities computationally derived from the drug-target interaction matrix to form the graph laplacian terms:

- Cosine similarity: computes the cosine of the angle between the two drug or target vectors. It is given by:

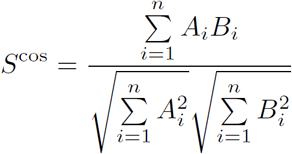
- Correlation: finds the the linear correlation (Pearson’s coefficient) between the drugs/targets. Its value ranges between −1 (denoting negative linear correlation) and +1 (denoting positive linear correlation). For two drug/target vectors (A and B) with sample size *n*, the correlation is calculated as follows:

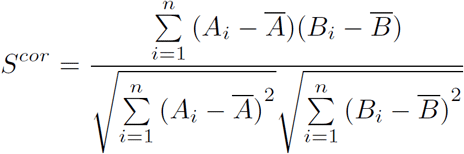

where

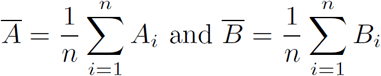
- Hamming similarity: is calculated by finding the complementary of hamming distance (percent of interaction positions differing between two drugs/targets) i.e. subtracting it from 1. It can be calculated as follows:

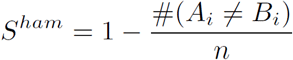
- Jaccard similarity: is defined as the percent of common interaction positions which are not zero for the two given pairs of drugs/target.

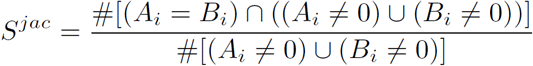

Table 1 gives a short summary of all the datasets.

**Table 1.** A summary of the number of Interactions, Drugs and Target in each dataset used.

| Datasets | Nuclear Receptor (NR) | G-Protein Coupled Receptor (GPCR) | Ion channel (IC) | Enzyme (E) |
| --- | --- | --- | --- | --- |
| # Interactions | 90 | 635 | 1476 | 2926 |
| # Drugs | 54 | 223 | 210 | 445 |
| # Targets | 26 | 95 | 204 | 664 |

### Pre-processing

As a first step, we sum up all the similarity matrices for drugs as well as targets and denote the combined similarity matrices by 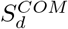 and 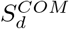 (equation (16)) respectively. Next, we sparsify them by keeping only the p-nearest neighbor of each drug/target profile. This sparsification, as shown by [24] helps the algorithm in learning a data manifold on or near to which the data is assumed to lie, preserving the local geometries of the original interaction data. Mathematically,

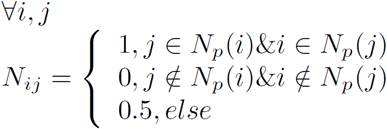

where *Np*(*i*) is the set of *p* nearest neighbors to drug *d*_*i*_. Similarity matrix sparsification is done by element-wise multiplying it with *N*_*ij*_. As a last pre-processing step, we compute the combined graph Laplacians. Also, since normalized version of graph lapacians are known to perform better [57], we use normalized graph laplacians 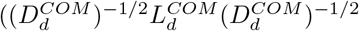 and 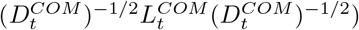 instead of graph laplacians (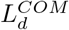 and 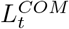).

### Evaluation

To evaluate our algorithm, we perform 5 repetitions of 10-fold cross validation and report the AUC and AUPR values for the test-interactions in the following 3 cross-validation settings: *cv-A*, *cv-B* and *cv-C*. *cv-A* randomly hides DTI pairs in the interaction matrix for testing/prediction; while *cv-B* and *cv-C* leave out the complete drug and target profiles respectively. The latter two cross-validation settings test the ability of the algorithm to perform when the drugs (in case of *cv-B*) and targets (in case of *cv-C*) are novel i.e. no interaction information is known int the interaction matrix.

We compare MGRDMF with some recent Matrix factorization based techniques proposed for DTI prediction under all the above mentioned croos-validation settings. These methods are very recent in the field and have already been compared against older methods [1]. Apart from the baselines (CMF [58] and GRMF [24]), the comparison of MGRDMF has also been done against a variant of GRMF proposed by us: Multi-GRMF, to observe how the incorporation of multi-graph regularization affects its performance.

Of note, in biological drug discovery, AUPR is a practically more important metric since it penalizes high ranked false positive interactions much more than AUC. This is because those pairs would be biologically validated later in the drug discovery process. As can be seen, MGRDMF always shows an improvement over MGRMF in terms of AUPR, moreover, with a significant margin when comparing against GRMF and CMF. The results clearly show that the muliple-graph regularization component plays an inegral role in DTI preidction improvement.

### Results

Our method, MGRDMF outperforms all the baseline techniques under the three cross-validation settings, as shown by the AUPR values in Tables 2, 4 and 6 and AUC values in Tables 3, 5 and 7. We performed cross validation on the training set to tune the parameters *p*, *μ*_1_, *μ*_2_, *k*_1_, *k*_2_ of our algorithm and hence find the best parameter combination for every dataset.

**Table 2.** Table showing AUPR values for DTI prediction with validation setting *cv-A*.

| AUPR | MGRDMF | MGRMF | GRMF | CMF |
| --- | --- | --- | --- | --- |
| E | **0.9746(0.0010)** | 0.9303(0.0015) | 0.8768(0.0020) | 0.8837(0.0026) |
| IC | **0.9652(0.0020)** | 0.9518(0.0005) | 0.9225(0.0022) | 0.9373(0.0019) |
| GPCR | **0.8345(0.0035)** | 0.8080(0.0031) | 0.7370(0.0024) | 0.7543(0.0017) |
| NR | **0.9175(0.0151)** | 0.7835(0.0225) | 0.6016(0.0378) | 0.6383(0.0149) |

**Table 3.** Table showing AUC values for DTI prediction with validation setting *cv-A*.

| AUC | MGRDMF | MGRMF | GRMF | CMF |
| --- | --- | --- | --- | --- |
| E | **0.9933(0.0007)** | 0.9906(0.0008) | 0.9647(0.0013) | 0.9705(0.0013) |
| IC | 0.9907(0.0008) | **0.9917(0.0014)** | 0.9747(0.0022) | 0.9832(0.0008) |
| GPCR | 0.9739(0.0013) | **0.9753(0.0022)** | 0.9432(0.0010) | 0.9493(0.0031) |
| NR | **0.9740(0.0096)** | 0.9573(0.0062) | 0.8892(0.0153) | 0.8679(0.0124) |

**Table 4.**
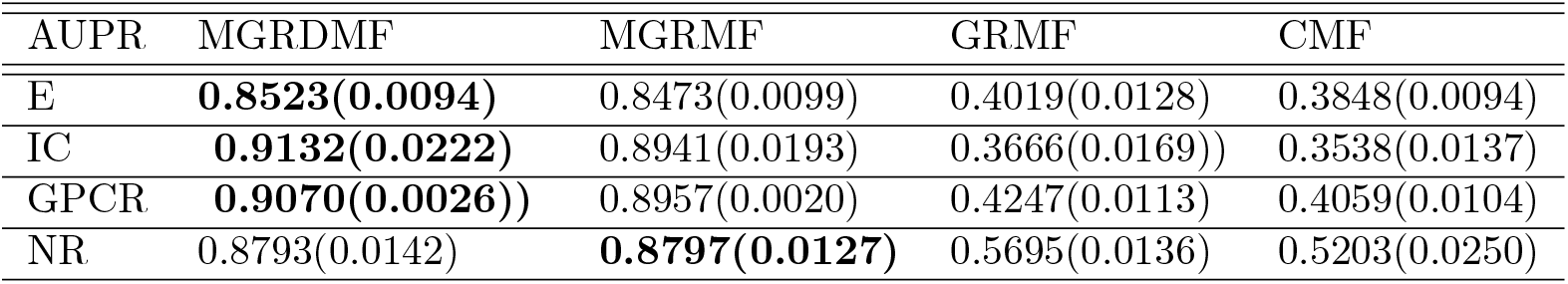
Table showing AUPR values for DTI prediction with validation setting *cv-B*.

**Table 5.** Table showing AUC values for DTI prediction with validation setting *cv-B*.

| AUC | MGRDMF | MGRMF | GRMF | CMF |
| --- | --- | --- | --- | --- |
| E | **0.9682(0.0028)** | 0.9456(0.0030) | 0.7982(0.0144) | 0.7952(0.0110) |
| IC | **0.9832(0.0126)** | 0.9668(0.0111) | 0.7902(0.0149) | 0.7576(0.0125) |
| GPCR | **0.9860(0.0008)** | 0.9733(0.0015) | 0.8800(0.0025) | 0.8067(0.0067) |
| NR | **0.9566(0.0109)** | 0.9474(0.0160) | 0.8615(0.0244) | 0.8124(0.0228) |

**Table 6.** Table showing AUPR values for DTI prediction with validation setting *cv-C*.

| AUPR | MGRDMF | MGRMF | GRMF | CMF |
| --- | --- | --- | --- | --- |
| E | **0.9237(0.0092)** | 0.9130(0.0127) | 0.8070(0.0185) | 0.7808(0.0131) |
| IC | **0.9142(0.0047)** | 0.9099(0.0035) | 0.8128(0.0069) | 0.7786(0.0108) |
| GPCR | **0.7485(0.0193)** | 0.7376(0.0291) | 0.6093(0.0134) | 0.5989(0.0323) |
| NR | **0.5735(0.0298)** | 0.5579(.0310) | 0.4643(.0183) | 0.4774(0.0173) |

**Table 7.**
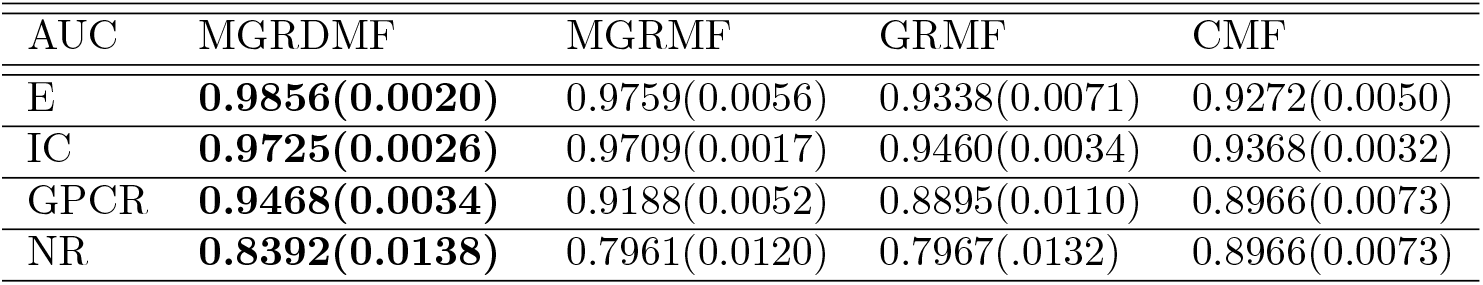
Table showing AUC values for DTI prediction with validation setting *cv-C*.

### Conclusion

In this paper, we introduce a novel computational approach, namely Multi-Graph Regularized Deep Matrix factorization (MGRDMF), to predict potential interactions between drugs and targets. The novelty of the proposed technique lies in structuring/regularizing drug-target interactions by multiple similarities of drugs and targets in the deep matrix factorization framework. The method is motivated by the success of deep learning and graph regularization in areas like content-based filtering, dimensionality reduction [59,60], clustering [61,62], semi-supervised learning [57], etc. It takes into account multiple Graph Laplacians over the drugs and targets for an improved interaction prediction.

We carried out the evaluation process using three cross-validation settings, namely *cv-A* (random drug-target pairs left out), *cv-B* (entire drug profile left out) and *cv-C* (entire target profile left out) for carrying out the comparison with three other state-of-the-art methods (specifically designed for DTI prediction). In almost all of the test cases, our algorithm shows the best performance, outperforming the baselines. One of the extensions of our work can be the incorporation of more/other types of similarity metrics for drugs/targets which could be either chemically or biologically driven or obtained from the metadata itself for improvement in the prediction accuracy. The proposed algorithm can also be used for interaction prediction in other bioinformatics problems such as protein-protein interaction [63], RNA-RNA interaction [64], etc. Not only this, it can be applied to other research areas such as collaborative filtering where multi-modality of the data can be captured using available metadata by finding the similarities for users and movies/items using user metadata (age, gender, occupation, etc) and item metadata (genre, year of release, etc).

